# Simulation study of factors affecting the accuracy of transcriptome models under complex environments

**DOI:** 10.1101/2024.08.30.610453

**Authors:** Dan Eiju, Yoichi Hashida, Taro Maeda, Atsushi J. Nagano

**Affiliations:** Graduate School of Media and Governance, Keio University, Fujisawa, Kanagawa 252-0882, Japan; Faculty of Agriculture, Takasaki University of Health and Welfare, Takasaki, Gunma, 370-0033, Japan; Institute for Advanced Biosciences, Keio University, Tsuruoka, Yamagata, 997-0017, Japan; Faculty of Agriculture, Ryukoku University, Otsu, Shiga, 520-2194, Japan

**Keywords:** Rice, Statistical modeling, Transcriptome, Environmental response Microarray, RNA-Seq

## Abstract

Characterization of molecular responses in real and complex field environments is essential for understanding the environmental response of plants. Field transcriptomics, i.e. modelling large amounts of transcriptomic and meteorological data, is the most comprehensive method of studying gene expression dynamics in complex environments. However, it is not clear what factors influence the accuracy of field transcriptome models. In this study, a novel simulation system was developed. Using the system, we performed a large-scale simulation to reveal the factors affecting the accuracy of the models. We found that the factors that had the greatest impact on the accuracy are, in order of importance, the expression pattern of the gene, the number of samples in the training data, the diurnal coverage of the training data, and the temperature coverage of the training data. Validation using actually measured transcriptome data showed similar results to the simulations. Our simulation system and the analysis results will be helpful for developing efficient sampling strategies for training data and for generating simulated data for benchmarking new modelling methods. It will also be valuable to dissect the relative importance of various factors behind transcriptome dynamics in the real environment.

**Key message:** Novel simulation system revealed how prediction accuracy of field transcriptome was affected by number and diversity of training data

## Introduction

In both natural and agricultural environments, plants are exposed to various fluctuating environmental factors, such as temperature, light, and rainfall. Plants are greatly affected by these environmental factors in various aspects, including growth and reproduction (Jones 2014; Taiz et al. 2023). The effects of environmental factors on crops are also large, and weather information is frequently used to predict crop yields (Chu and Yu 2020; de Los Campos et al. 2020; van Klompenburg et al. 2020). Plants adapt to their environment through changes in gene expression, metabolism, and morphology. Plant responses to the environment have been widely studied at the molecular level. Most of them are conducted in simple and limited laboratory environments. However, plants live in field environments where various factors fluctuate intensely. Therefore, to understand the reality of plant responses to the environment, it is important to clarify the responses of plants to complex environmental stimuli in field environments at the molecular level (Poorter et al. 2016; Annunziata et al. 2017; Matsubara 2018; Song et al. 2018; Dantas et al. 2021; Hashida et al. 2022)

Gene expression analysis is one of the most widely used methods for investigating the environmental response of organisms at the molecular level (Cerda and Alvarez 2024; Kim et al. 2024; Cai et al. 2024). Transcriptome analysis is a method for comprehensively quantifying gene expression, and there are numerous examples of its use in laboratory research (Yu et al. 2022; Lim et al. 2022). However, there have only been a limited number of studies that have conducted gene expression analysis or transcriptome analysis in field environments (Andersson et al. 2004; Sjödin et al. 2008; Aikawa et al. 2010; Hayes et al. 2010; Sato et al. 2011; Levy et al. 2011; Izawa et al. 2011; Richards et al. 2012; Nagano et al. 2012; Satake et al. 2013).In recent years, there are growing interest in the differences between the field and the laboratory (Hashida et al. 2022). Along with this, studies of field transcriptomes are also increasing. Studies have been reported not only on crops such as rice (Plessis et al. 2015; Groen et al. 2020; Matsunami et al. 2023; Ushio et al. 2023) and barley (Hirayama et al. 2020), but also on wild plants such as *Arabidopsis halleri* (Nagano et al. 2019; Komoto et al. 2024; Yumoto et al. 2024), woody plants (Nose and Watanabe 2014; Fasoli et al. 2018; Satake et al. 2019; Nose et al. 2020; Lu et al. 2020; Shirasawa et al. 2022; Nose et al. 2023; Tornielli et al. 2023; Lihavainen et al. 2023; Satake et al. 2023; Suhaimi et al. 2023), and animals (Nakayama et al. 2023).

The FIT model of rice is one of the most comprehensive studies to date that has modeled the relationship between meteorological factors and gene expressions in plants growing in the field (Nagano et al. 2012). In developing the FIT model, the leaves of rice plants grown in paddy fields were sampled diurnally on multiple days throughout the entire cultivation period, and the transcriptomes of hundreds of samples were measured. Regression models were then trained to predict rice gene expression using meteorological data (air temperature, relative humidity, global solar radiation, atmospheric pressure, wind speed, and precipitation), circadian clock and plant age as explanatory variables. Regression models were obtained for 96.7% of the genes expressed in rice leaves. For example, the model for the *OsGI* gene, which is a component of the circadian clock, reproduced the clear diurnal rhythm and the change in amplitude dependent on nighttime temperature. The FIT model made it possible to obtain a computational prediction of the expression level of each gene under any given weather condition (Iwayama et al. 2017). Furthermore, applications of the FIT model have been developed, such as quantitative genetic analysis of gene expression dynamics in the field (Kashima et al. 2021) and improving generalization performance by combining systematic measurements in diverse controlled environments (Hashida et al. 2024). However, the difficulty in obtaining enough data for constructing the model is an obstacle to its widespread use in a wide variety of species and cultivars. In general, the more training data used, the higher the accuracy of the model is expected to be, but simultaneously the cost of acquiring the data increases. To successfully handle this trade-off, it would be useful to clarify the impact of the number of training data and the diversity of environmental conditions on the prediction accuracy.

In this study, we constructed an *in silico* simulation system for field transcriptome prediction and examined the effects of the number and diversity of training data. Through large-scale simulations, we were able to quantitatively clarify the magnitude of the impact of various factors on prediction accuracy. The simulation system and analysis results in this study will advance our understanding and prediction of environmental responses in the field.

## Materials and Methods

### Overview of Simulation

Simulation in this study is divided into three major steps: 1) Preparation of reference FIT model, 2) Development of test FIT models, and 3) Evaluation of model accuracy (Fig. 1a). In the “1) Preparation of reference FIT model,” reference FIT model was created using the R package FIT (Iwayama et al. 2017) with gene expression levels measured on 461 actual field samples and correspond weather data as input (Nagano et al. 2012). Next, in the “2) Development of test FIT models”, the sampling conditions to be verified were input into the reference FIT model, and based on the output gene expression data, test FIT models were created. In the “3) Evaluation of model accuracy”, the test FIT models (494 conditions) and the reference FIT model were given the same weather conditions for validation, and the test FIT models were evaluated by comparing their predictive gene expressions (108 time-point).

**Figure 1.**
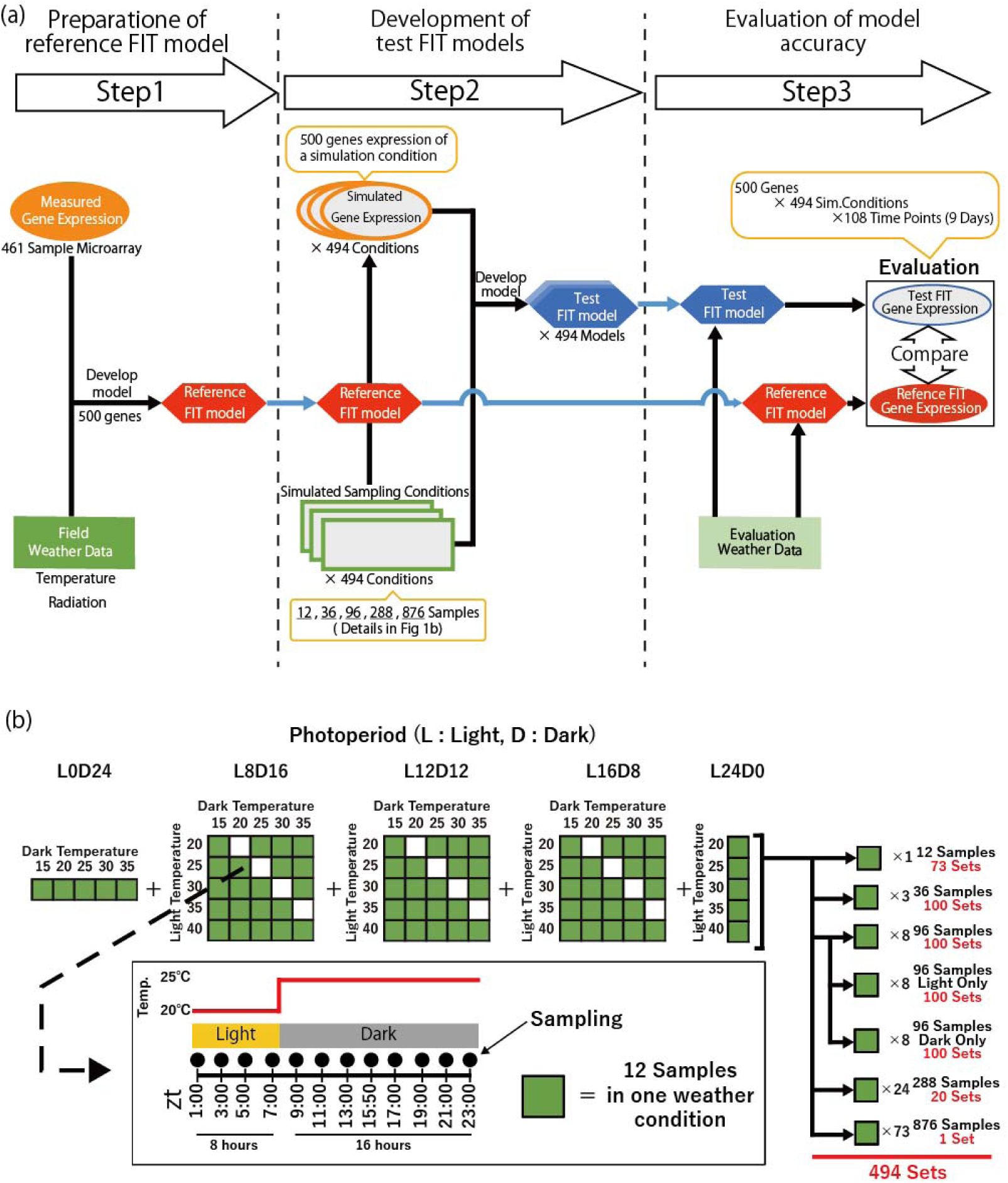
Overview of the simulation. (**a**) Flowchart of the simulation. The simulation consists of three steps. The circles indicate gene expression data. The hexagons indicate FIT models. The squares indicate weather data. (**b**) Details of the 494 simulation sampling conditions. Each green square represents one weather condition. A weather condition consists of a combination of light temperature, dark temperature, and photoperiod. In one weather condition, 12 samples are obtained at two-hour intervals, as shown in the examples in the lower half of this panel. Simulation sampling conditions consist of 1, 3, 8, 24 or 73 weather conditions.

### Preparation of reference FIT model (Step1)

To create the reference FIT model, we used actual transcriptome data of field samples and weather data in our previous study (Nagano et al. 2012). In the previous study, rice seeds were sown in germination boxes and transplanted to paddy field at 30 days or 14-15 days after germination. 461 samples of the youngest fully expanded leaves were collected for DNA microarray analysis. The weather data were obtained from the Tateno weather station of Japan Meteorological Agency located near the field in Tsukuba, Japan (36°02′N, 140°10′E, altitude 25 m asl). In this study, we used air temperature (°C) and global solar radiation (kJ m-2min-1) as input data to train the reference FIT model. To reduce the computational cost, we performed affinity propagation of genes based on the measured transcriptome data and selected 500 genes representing each cluster for analysis in this study. The 500 genes maintained the diverse expression pattern of the entire transcriptome (Supplementary Table S1).

### Development of test FIT models (Step2)

Simulation sampling conditions were created by combining 73 different weather conditions (Supplementary Table S2) that could be reproduced in conventional growth chambers. A total of 494 simulation sampling conditions were created with various numbers of weather conditions (1, 3, 8, 24, and 73) (Fig.1b, Supplementary Tables S3∼S7). The 73 weather conditions were combinations of day length (L0D24, L8D16, L12D12, L16D8, L24D0, representing 0, 8, 12, 16, 24 h light periods, respectively) and light temperature (20, 25, 30, 35, 40°C) and dark temperature (15, 20, 25, 30, 35°C), excluding conditions with the same temperature during the light and dark periods (Supplementary Table S2). It was assumed that 12 time points would be sampled under 1 weather condition. The simulation would have 36 samples for 3 weather conditions, 96 samples for 8 weather conditions, 288 samples for 24 weather conditions, and 876 samples for 73 weather conditions. The expected gene expression for each weather condition was calculated using the reference FIT model, assuming that transcriptome analysis was conducted at 12 time points: every 2 hours from 01:00 to 23:00 on 10 days after transplanting. To simulate measurement error by microarray, we added random values following normal distribution to the expected gene expressions. The standard deviation (SD) of the regression residuals for each gene in the step 1 was used as SD of the normal distribution for simulated measurement error. The simulated gene expressions with measurement error were used as training data for developing test FIT models. We successfully obtained the test FIT models with optimized parameters except in 5 conditions with L24D0 of 12 samples and 5 conditions with L0D24 of 12 samples. In the light-only condition, two samples were sampled at 1, 3, 5, 7, 9, 11 h after the start of the light period, and in the dark-only condition, two samples were sampled at 13, 15, 17, 19, 21, 23 h after the start of the light period.

### Evaluation of model accuracy (Step3)

To evaluate the test FIT models developed from 494 simulation sampling conditions, the same evaluation weather data were input into both the reference FIT model and each test FIT model, and gene expressions predicted by the reference and test FIT models were then compared. Extreme gene expression values from the test FIT model, representing 3.4% of the data, were treated as outliers. Values below 0 were set to 0, and those above 20 were set to 20 for analysis.

Pearson product moment correlation coefficients and root mean squared error (RMSE) were calculated from the gene expressions of the reference and test FIT models for each day of the evaluation weather data. There were cases where the variance of these values was zero and correlation coefficients could not be defined, which was the case for 5.5% (120601/2178000) of the total and was excluded from the analysis.

The weather data from May 1, 2008, to October 31, 2008 in the Tateno weather station of Japan Meteorological Agency ware used as the evaluation weather data. We assumed that the transcriptomes were obtained at 12 time points: every 2 hours from 01:00 to 23:00, on 9 days: every 2 weeks from June 7, 2008, to September 27, 2008.

### RNA-Seq data of rice plants actually cultivated in 73 conditions

To evaluate our simulation results, we used actual RNA-Seq data reported in our previous study (Hashida et al. 2024). The RNA-Seq data were measured from rice plants cultivated in high irradiance growth chambers (GC, ECP101, TECS, Ibaraki, Japan) with 73 different conditions corresponding 73 weather conditions in this study (Supplementary Table S2). Under each condition, sampling was then conducted every 3 h for 24 h (1.5, 4.5, 7.5, 10.5, 13.5, 16.5, and 19.5h after the acclimation period, 8 times in total). The uppermost, fully expanded leaves were sampled from one plant per sampling point. Further details of the experimental procedures of RNA-Seq were described in Hashida et al. 2024.

## Result

### Prediction accuracy improved with increasing sample size

The impact of different sample sizes used for training on the prediction accuracy of the test FIT model was evaluated. For each gene, we calculated the correlation coefficient and RMSE for predicted gene expression levels (at 108 time points) by the reference and test FIT models. The mean values of the correlation coefficient and RMSE for 500 genes were calculated for each simulation sampling condition (Fig. 2a, b). The results showed that both the correlation coefficient and RMSE improved as the sample size increased (P < 0.01, Mann-Whitney U test). For example, the median correlation coefficient in the simulation condition with 12 samples was 0.27 but improved to 0.64 when the sample size increased to 288. The RMSE similarly improved from 2.77 to 1.03. While no significant difference was noted when the sample size further increased from 288 to 876, the mean value showed a slight improvement.

**Figure 2.**
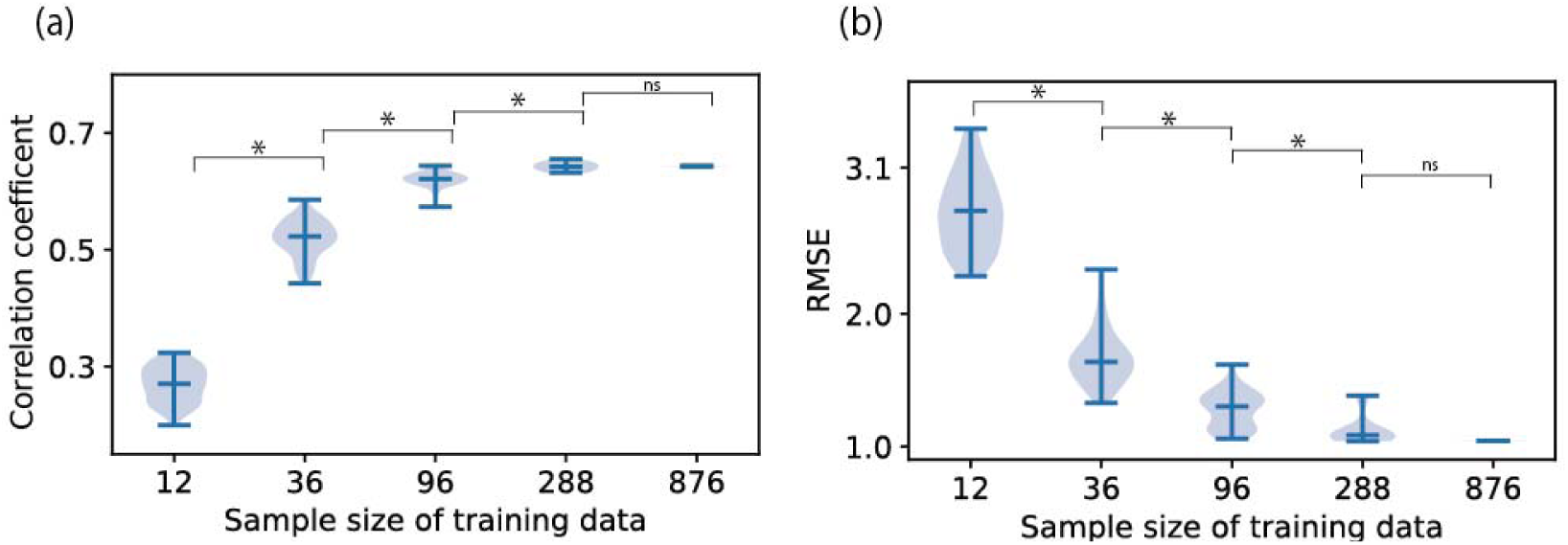
Prediction accuracy on different sample size. (**a, b**) Correlation coefficient (**a**) and RMSE (**b**) on different sample size of training data. The horizontal line in the violin plot is the median. For each of the 500 genes in each simulation sampling conditions, we calculated Pearson’s correlation coefficient and RMSE between the test and reference FIT model. The correlation coefficients and RMSE for the 500 genes were averaged in each simulation sampling condition. n = 73, 100, 100, 20, 1 for 12, 36, 96, 288, 876 samples of training data, respectively. *: P < 0.001. ns: not significant.

### Prediction accuracy depended on date of evaluation data

To examine further details of the simulation results, we focused on the temporal gene expression pattern of each gene under each simulation sampling condition. As an example, the results of three typical genes under the same simulation sampling condition were shown (Fig. 3a-c). In addition to genes with good/poor prediction accuracy on all days of evaluation data (Fig. 3a and b), there were also a gene with significantly different prediction accuracy depending on the day of evaluation data (Fig. 3c). In the example shown in Fig. 3c, the prediction was successful in the early period of the time series. However, in the later period, although the fluctuation pattern of gene expression levels could be predicted, the average expression levels could not be predicted. Examples where the prediction accuracy varies greatly depending on the day of the evaluation data motivated us to evaluate the prediction accuracy for each day for each individual gene. We showed the distribution of the correlation coefficient and RMSE for each day of evaluation data for each individual gene (Fig. 3d).

**Figure 3.**
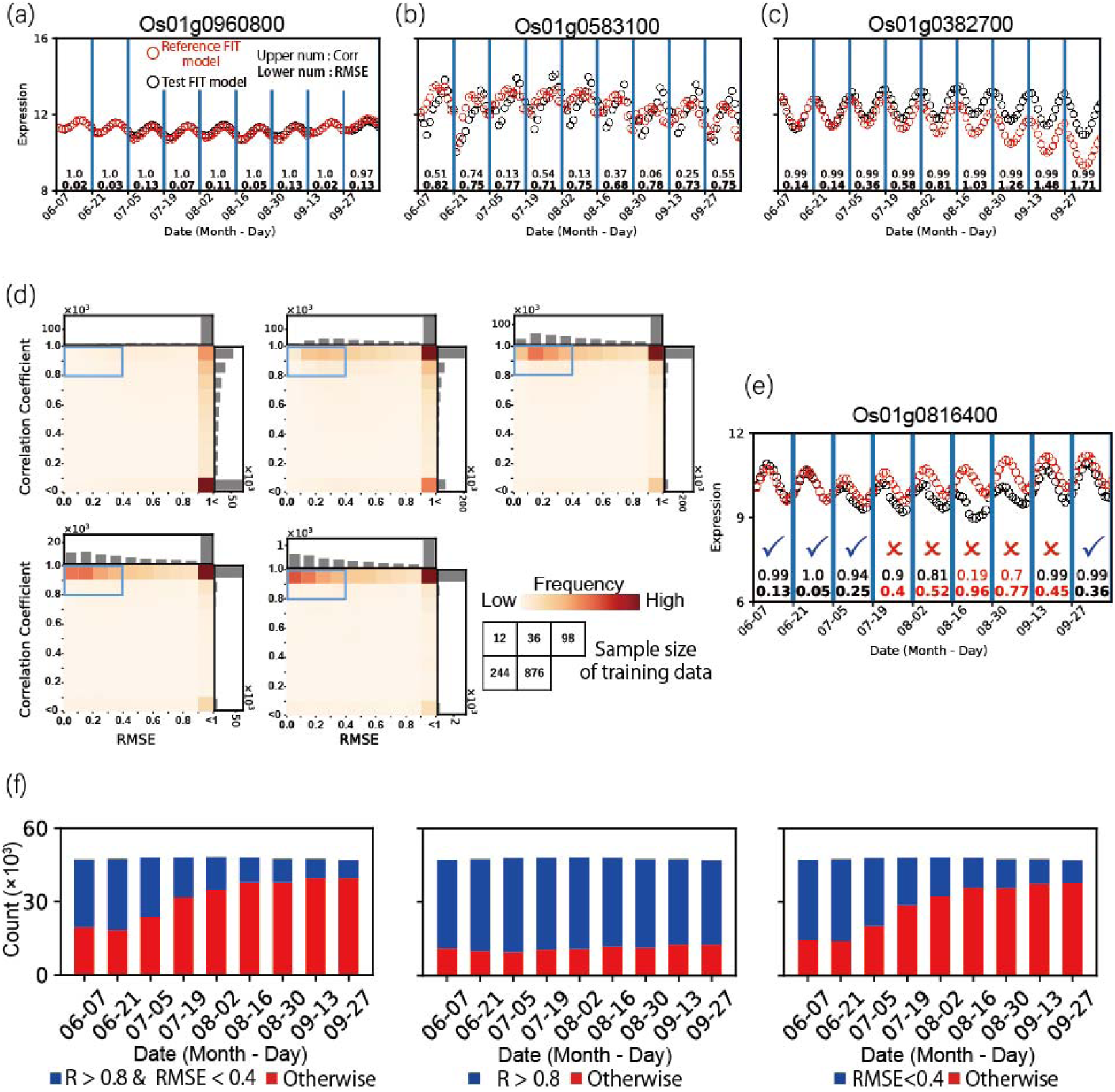
Evaluation of prediction for each day of evaluation data. (**a-c**) Examples of time series. Genes with good (**a**) and poor (**b**) prediction accuracy. A gene with different prediction accuracy depending on the day of evaluation data (**c**). Red and black circles indicate gene expressions by the reference and test FIT model, respectively. The test FIT models of these 3 genes were trained in the same simulation sampling condition. Correlation coefficient and RMSE of each day are indicated at the bottom of the plot. (**d**) Frequency of correlation coefficient and RMSE in various sample sizes of training data. The blue box indicates the threshold for determining a successful prediction: correlation coefficient > 0.8 and RMSE < 0.4. n = 260757, 425098, 428405, 86527, 4342 for 12, 36, 96, 288, 876 samples of training data, respectively. (**e**) An example of the results of the predictions being classified as successful (check mark) or unsuccessful (cross mark). From July 19 to September 13, the predictions were unsuccessful, but on other days they were successful. (**f**) Counts of data exceeding the threshold for the correlation coefficient (R) and RMSE in sample size 96. The correlation coefficient and RMSE were calculated for each gene, each simulation sampling condition, and each day of the evaluation data.

To simplify the analysis, we decided to classify the prediction results into two categories: successful or unsuccessful, because we had to treated 2,223,000 prediction results that were consistent with combinations of 9 days of evaluation data, 494 simulation conditions and 500 genes. In this study, we set the threshold for determining successful or unsuccessful predictions at “correlation coefficient > 0.8 and RMSE < 0.4” (Fig. 3d, blue box). 19.7% of the entire data set had correlation coefficient more than 0.8 and RMSE less than 0.4. Using this threshold, for example, for the Os01g0816400 gene under a simulation condition with 96 samples, the prediction was unsuccessful from June19 to September13, but successful on other days (Fig. 3e). When we used “correlation coefficient > 0.8” and “RMSE < 0.4” as individual criteria, the trends of successful predictions on days of evaluation data were different between correlation coefficient and RMSE (Fig. 3f, Supplementary Fig. S1). In correlation coefficients, the rates of successful prediction were similar among days of evaluation data. In RSEM, the rates of successful prediction gradually decreased toward the later days of evaluation data.

### Prediction accuracy depended on simulation sampling conditions and gene expression patterns

To investigate the impact of simulation sampling conditions and gene on prediction accuracy, we show how many days of the nine-day evaluation data were successfully predicted (Fig. 4a). When focusing on the simulation sampling conditions, there was no major difference in the number of genes that were successfully predicted, although there were slightly more genes that were successfully predicted under some conditions. There was a tendency for the successfully predicted genes to increase as the samples increased.

**Figure 4.**
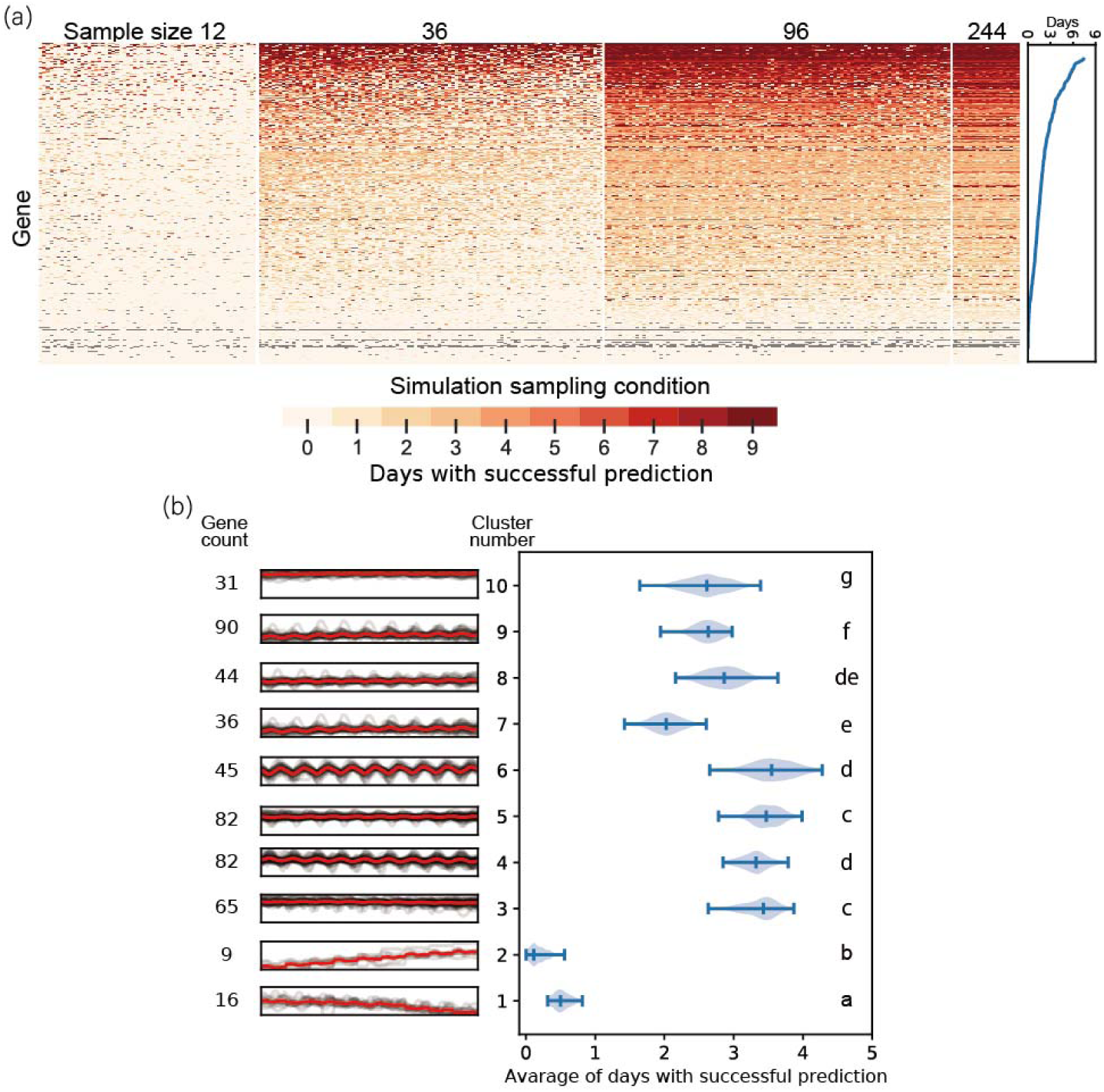
Prediction accuracy depends on genes and simulation sampling conditions. (**a**) Days with successful prediction of each gene for each sampling condition. The genes on the vertical axis are ordered by mean of days with successful prediction in all simulation sampling condition. The simulation sampling conditions on the horizontal axis are ordered by mean of days with successful prediction by each sample size. The plot on the right-side shows means of days with successful prediction in simulation sampling conditions with 96 samples. (**b**) Prediction accuracy for various gene expression patterns. The temporal patterns of gene expressions from the reference FIT model of 500 genes are clustered (left). Gray lines indicate gene expressions of each gene. Red line indicates mean expressions of genes in a cluster. The average numbers of days with successful prediction of genes in the clusters (right). Different letters indicate significant differences (*: P < 0.001., Steele-Duwas test)

On the other hand, when focusing on genes, it was found that the number of simulation sampling conditions with successful predictions varied greatly depending on the gene. There were genes that were successfully or unsuccessfully predicted in almost all simulation sampling conditions (Fig. 4a). There were also genes that were only successfully predicted under specific conditions, and interestingly, the conditions with successful predictions differed depending on the gene (Fig. 4a).

To investigate the factors behind the differences in prediction accuracy among genes, we examined the time-series expression patterns and gene expression noise among individuals. We clustered the temporal gene expression patterns of the reference FIT model and examined the average prediction accuracy of the test FIT models for genes in each cluster (Fig. 4b). We found that the average prediction accuracy differed significantly among clusters (Fig. 4b right, P < 0.001, Steele-Duwas test). In particular, the prediction accuracy was low for genes whose expression levels changed over the long term throughout the evaluation data period (cluster 1, 2). Next, we analyzed the effect of gene expression noise among individuals. In general, the variance of expression across biological replicates differs depending on the gene. In our simulation, when preparing the training data for the test FIT model, we added artificial noise corresponding to the magnitude of actual gene expression noise among individuals (see method for details). In genes with larger noise relative to their deterministic expression dynamics, prediction accuracy may be lower. Therefore, we analyzed the correlation between the coefficients of determination of the reference FIT models and the average number of successful prediction days of test FIT models (Supplementary Fig. S2). A slight correlation was observed, but the correlation decreased with increasing sample size.

### Diurnal sampling contributes precise prediction of diurnal expression oscillation

We analyzed the impact of diurnal sampling on prediction accuracy. Because of diurnal environmental factors such as temperature and light, as well as the circadian clock, field transcriptomes are expected to exhibit significant diurnal variation. For this reason, diurnal sampling is important. However, since circadian sampling requires a great deal of effort, it is occasionally limited to sampling during a narrow time period, such as only during the daytime. To clarify the impact of diurnal sampling on prediction accuracy, we compared the prediction accuracy of diurnal sampling conducted throughout the light and dark periods (hereafter, the control condition) with that of sampling conducted for only half a day in either the light or dark period (hereafter, the light-only condition and the dark-only condition). In the control condition, it was assumed that sampling would be carried out at 12 points per day, with one replicate. In the light-only and dark-only conditions, it was assumed that sampling would be carried out at 6 points per half-day in the light (or dark) period, with two replicates (Fig. 5a). Therefore, it was possible to compare the control condition with the light-only and dark-only conditions, with the number of samples being equal.

**Figure 5.**
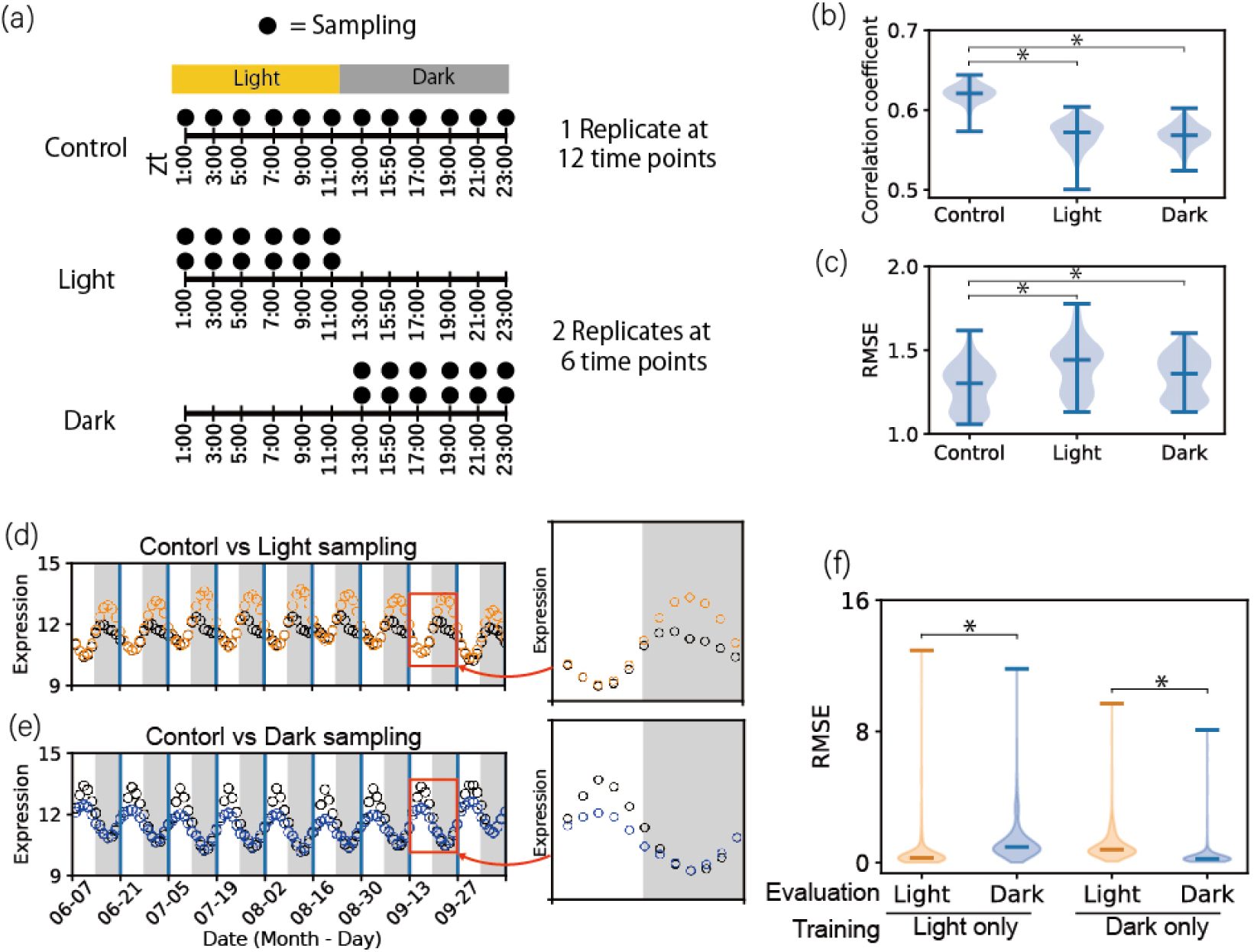
Effect of diurnal sampling on prediction accuracy. (**a**) Scheme of sampling schedules for the control, light-only and dark-only conditions. In the control condition, sampling is carried out at 12 points per day, with one replicate. In the light-only and dark-only conditions, sampling is carried out at 6 points per half-day in the light (or dark) period, with two replicates. (**b, c**) Correlation coefficient (**b**) and RMSE (**c**) on the control, light-only and dark-only conditions. The correlation coefficients and RMSEs for the 500 genes were averaged in each simulation sampling condition. n = 100 for all three conditions. *: P < 0.001. (**d, e**) Examples of genes that were unsuccessfully predicted in the light-only (**d**) or dark-only (**e**) conditions but were successfully predicted in the control conditions. Black, orange and blue circles indicate gene expressions of the control, light-only and dark-only conditions, respectively. Gray area indicates dark period. Plot on Sep. 13 is enlarged at right side. (**f**) RMSE during sampled/unsampled time periods. Focusing on genes with successful predictions for 9 days under the control conditions and for no day under the light-only or dark-only conditions. *: P < 0.001

To compare the prediction accuracy of the light/dark only condition with the control condition, the correlation coefficient and RMSE were calculated from the gene expression levels of the test FIT model for each gene, and the correlation coefficients and RMSE of 500 genes were averaged for each condition. As a result, the prediction accuracy of the control condition was the highest in terms of both the correlation coefficient and RMSE (Fig. 5b, c). To clarify the prediction accuracy during unsampled time periods in the light-only/dark-only conditions, we show the time-series expression plots of genes that failed to be predicted in the experimental sampling conditions but were successfully predicted in the control conditions (Fig. 5d, e, Supplementary Fig. S3a). Although the relative fluctuation patterns of gene expression could be predicted, the gene expression levels differed from the control conditions during the unsampled time periods (dark period in the case of sampling only during the light period) (Fig. 5d, e). When RMSE was calculated separately for the unsampled and sampled time periods, the RMSE value was greater during unsampled time periods (P < 0.01, Mann-Whitney U test) (Fig. 5f).

### Prediction accuracy in the unsampled temperature range

We examined the impact of the temperature of the simulation sampling conditions on prediction accuracy. For example, in some simulation conditions, the model is trained using data from 20°C to 30°C, but the evaluation data includes data below 20°C and above 30°C, so there is a possibility that the prediction accuracy will decrease in the temperature range that was not used to train the model (Fig.6a). We examined how much the temperature of the evaluation weather data was covered by the temperature of the training data for test FIT models (Fig.6b, Supplementary Fig. S4a). 47% of the evaluation weather data fell within the range of training data in the simulation sampling conditions with 96 samples. There was not much difference depending on the date of the evaluation data, but there were relatively few data outside of the range in the middle of the growth period.

**Figure 6.**
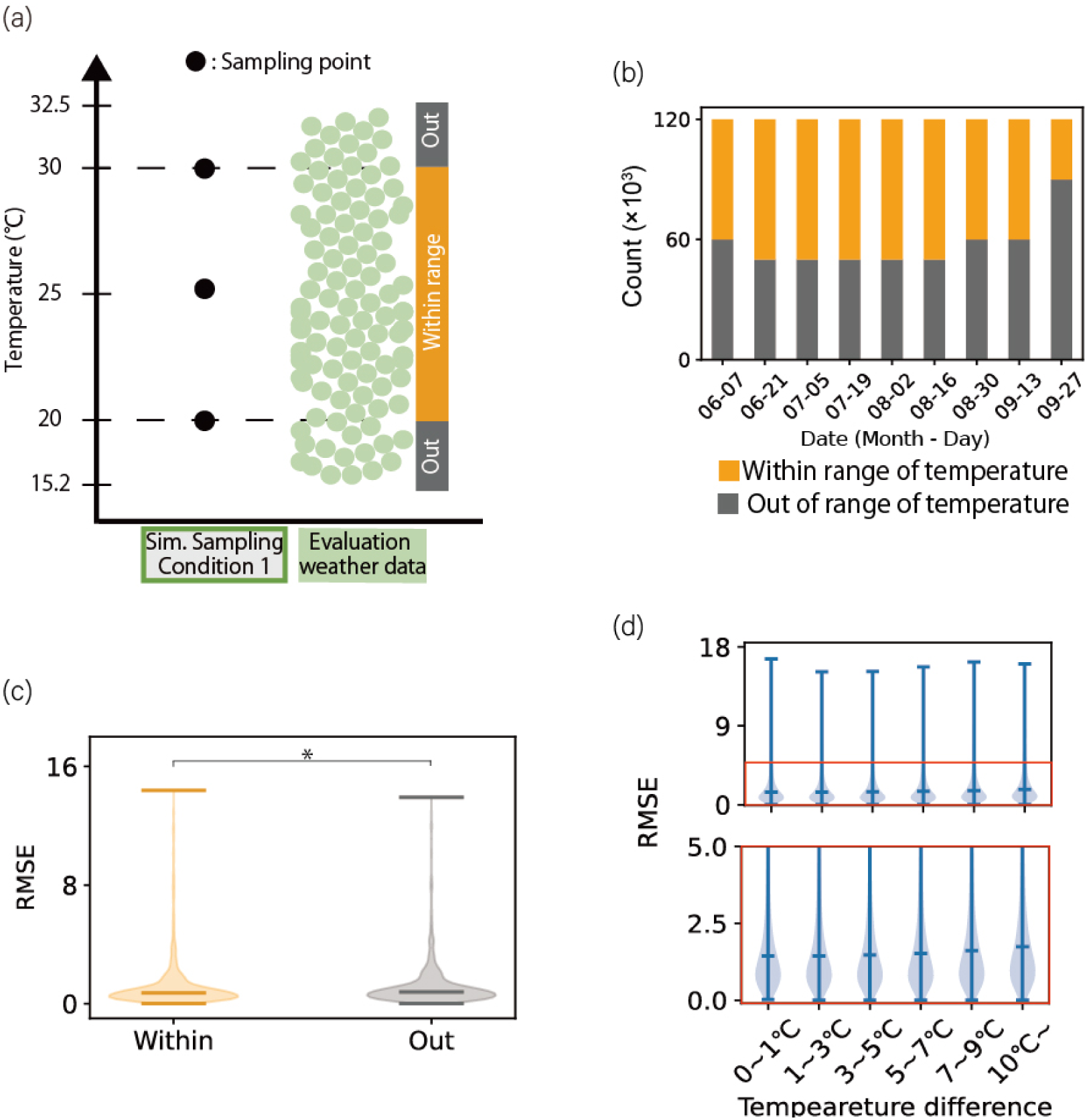
Prediction accuracy in the unsampled temperature range. (**a**) An example of temperature used in the training data and in the evaluation data. In this example, temperature of training data ranges from 20 to 30°C. Temperature of the evaluation data is distributed out of the range of temperature of training data. (**b**) Data counts temperature of the evaluation data within (orange) and out of (gray) the range of training data. (**c**) RMSE within and out of the temperature range of the training data, for simulation sampling conditions with 96 samples. *: P < 0.001 (**d**) RMSE out of the temperature range of training data with 12 samples. Area in the red box of the upper plot is enlarged as the bottom plot. Prediction errors are shown for each temperature difference between the temperature of the training data and that of the evaluation data. Prediction errors are different among groups (P < 0.001, Steele-Duwas test).

We investigated the relationship between the RMSE of gene expression and whether the temperatures of the evaluation data fell within the range of the training data (Fig. 6c). The RMSE for gene expression were significantly smaller within the range than outside of the range of the training data (P < 0.01, Mann-Whitney U test), but the difference was not large. We examined whether the RMSE of gene expression levels depended on the magnitude of the temperature difference between the training and evaluation data, using simulation sampling conditions with 12 samples consisting of only one temperature condition. As the temperature difference increased, RMSE significantly increased (P<0.001, Steele-Duwas test), but the extent of the increase was small (Fig. 6d). Overall, it was found that the effect of the temperature conditions of the training data on the prediction accuracy was limited.

### Validation of the simulation results with actual data

The simulation results were verified by analysis using actually measured gene expression data. Two approaches were used for the validation. The first is to train a reference FIT model using a subset of the measured data. The second is to train a FIT model using gene expression levels measured by RNA-seq from rice plants grown under conditions corresponding to the simulation sampling conditions.

In the first approach, we investigated the effects of light/dark conditions and temperature. 461 samples of microarray data used to train the reference FIT model were divided into two subsets: those with zero solar radiation at the time of sampling and those with non-zero solar radiation (Supplementary Fig. S5). We obtained two FIT models trained by the two subsets: the dark-only FIT model and the light-only FIT model. The predicted gene expressions of the reference FIT model for the evaluation weather data were compared with those of the dark-only/light-only FIT model. When RMSE was calculated separately for dark and light time points of the evaluation weather data, the error of the dark-only FIT model was greater for light time points than for dark time points, and the error of the light-only FIT model was the opposite (Fig. 7a, P < 0.01, Mann-Whitney U test). These results were consistent with the simulations (Fig. 5f). We extracted a subset with temperature from 20 to 30°C from the data used to train the reference FIT model. The temp-lim FIT model was trained by the subset. RMSE was calculated separately for the evaluation weather data within 20 to 30 °C and outside of the range. RMSE of the temp-lim FIT model was lower for evaluation data within the temperature range, 20 to 30 °C, than those outside of the range (Fig. 7b, P < 0.01, Mann-Whitney U test). This was consistent with the simulations (Fig. 6c).

**Figure 7.**
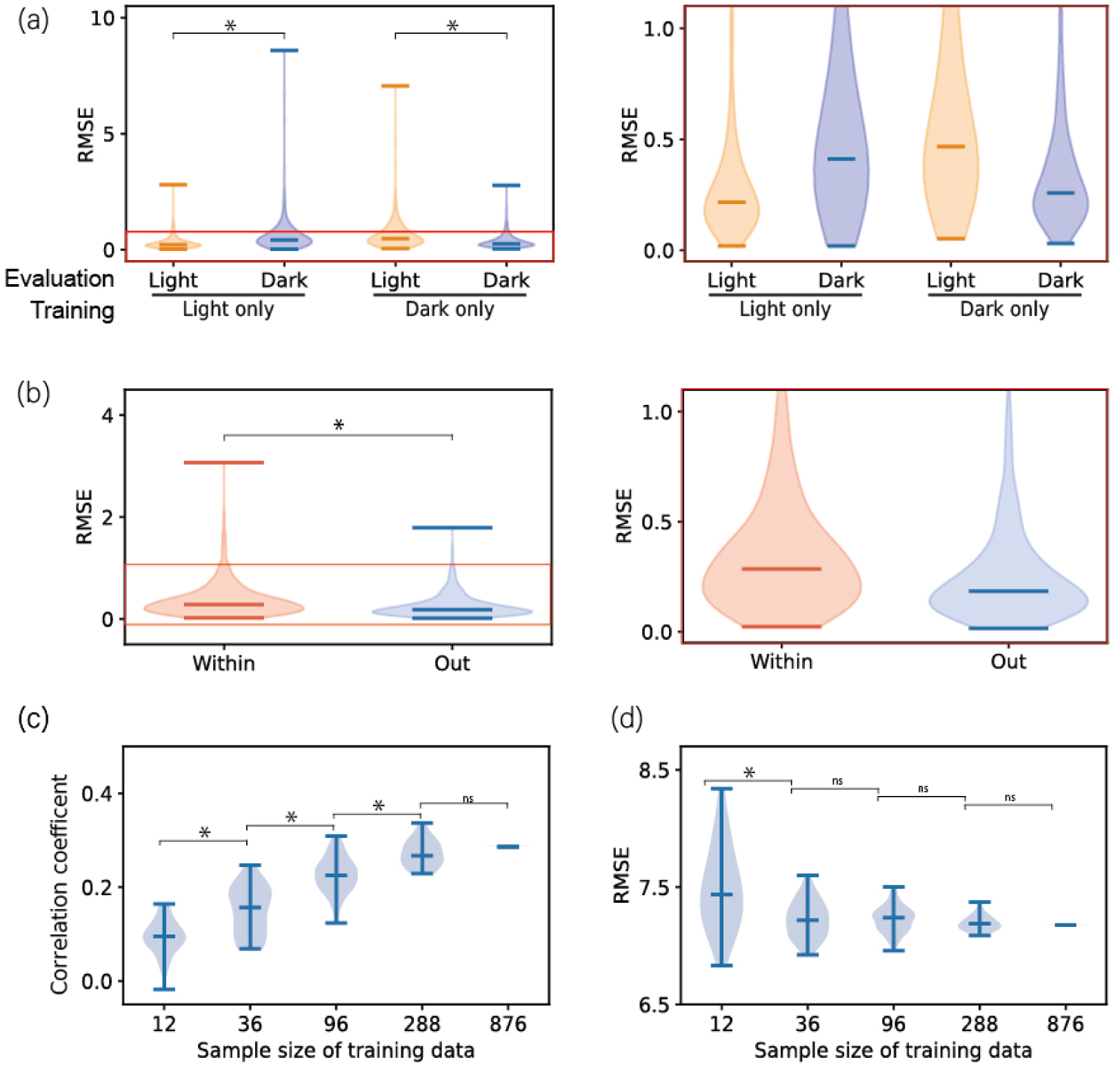
Validation of the simulation results with actual data. (**a**) Prediction error of the light-only/dark-only FIT model trained by subset of actually measured gene expression data. The prediction errors are summarized by light/dark periods of evaluations data. *: P < 0.001. Area in the red box of the left plot is enlarged as the right plot. (**b**) Prediction error of the temp-lim FIT model trained by a subset with temperature from 20 to 30 °C of actually measured gene expression data. The prediction errors are summarized by within or out of the temperature range of the training data. *: P < 0.001. Area in the red box of the left plot is enlarged as the right plot. (**c, d**) Correlation coefficient (**c**) and RMSE (**d**) between the reference FIT model and the FIT models trained by actual RNA-Seq data. The correlation coefficients and RMSEs for the 500 genes were averaged in each simulation sampling condition. *: P < 0.001. ns: not significant.

In the second approach, we used RNA-Seq data of rice plants that were actually grown in growth chambers corresponding to the 73 weather conditions of the simulations (Fig. 1b green squares) (Hashida Y, Kyogoku D, Tanaka SE, Mori N, Tanabata T, Watanabe H, Nagano AJ 2024) The actual transcriptome data, instead of the simulated gene expressions, were used to train the FIT model. The predicted expression were obtained by inputting the evaluation weather data into the FIT model trained by the measured data. The predicted expressions were compared with those of the reference FIT model. When we looked at the change in prediction accuracy with increasing sample size, we found that both the correlation coefficient and RMSE improved with increasing sample size (Fig. 7c, d, P < 0.01, Mann-Whitney U test).

## Discussion

This study revealed that the factors that have the greatest impact on the accuracy of field gene expression prediction are, in order of importance, expression pattern of the gene, the number of samples in the training data, the diurnal coverage of the training data, and the temperature coverage of the training data (Fig. 8). In genes whose expression levels changed depending on the growth stage, the prediction accuracy was relatively low (Fig.4b). In the simulations of this study, data on the gene expression levels in the late growth stage were not included for training the model. This bias in the training data is thought to be the cause of the low prediction accuracy for these genes. To improve the prediction accuracy for these genes, it might be effective to include data from various growth stages during training. As the number of samples in the training data increased, the prediction accuracy improved, but the degree of improvement gradually decreased (Fig. 4a). Under the simulation sampling conditions of this study, it appeared that the prediction accuracy saturated at several hundred samples. However, the number of samples at which accuracy saturates may be underestimated, because actual field measurements are affected by fluctuations of minor factors that are not included in the simulation system(Hashida et al. 2022; Matsunami et al. 2023; Ushio et al. 2023). The decrease in prediction accuracy was relatively small at temperatures outside the training data (Fig. 6b). On the other hand, in the case of sampling for half a day (only during the light and dark periods), the prediction accuracy decreased considerably during the time periods that were not sampled (Fig. 5f). The cause of this decrease could be the circadian clock, light conditions, or both, but it is not possible to conclude which is the cause in this simulation.

**Figure 8.**
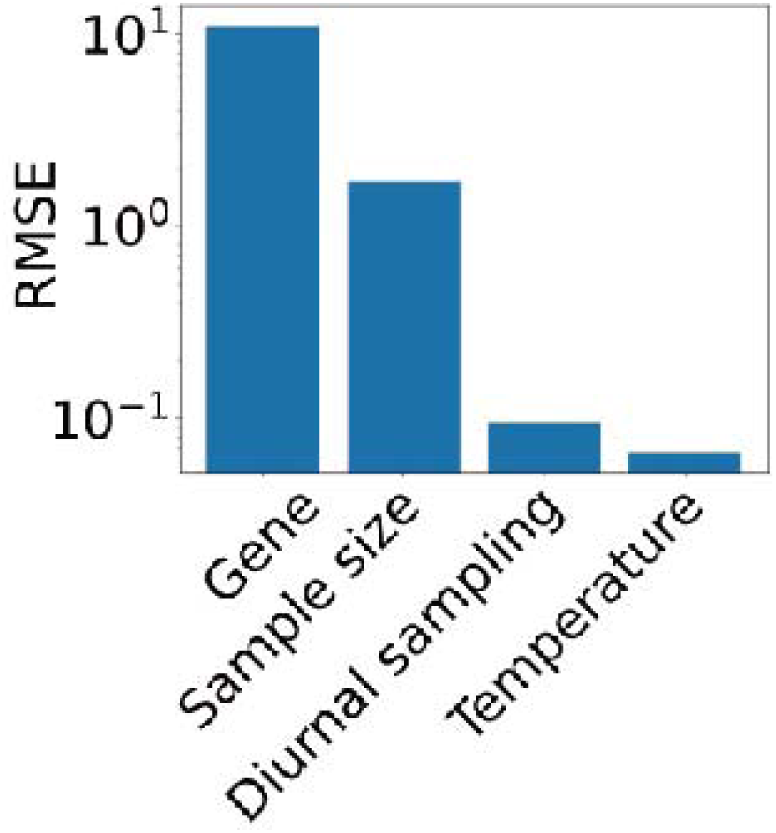
The impact of various factors on the accuracy of gene expression prediction. Gene: the difference between average RMSE values of the best predicted gene and the worst predicted gene (Fig.4a). Sample size: the difference between average RMSE values for the training data with 876 samples and for those with 12 samples (Fig.2b). Diurnal sampling: the difference between average RMSE values for the control condition and the light-only / dark-only conditions (Fig.5c). Temperature: the difference between average RMSE values within and out of the temperature range of training data (Fig.6c).

The simulations in this study assumed the gene expression levels measured by microarrays. This is because the most well-validated field transcriptome data was obtained using microarrays, and this study used that data. In recent years, RNA-Seq has become the primary method for transcriptome analysis due to the development of various high-throughput RNA-Seq library preparation methods (Adiconis et al. 2013; Kamitani et al. 2019; Janjic et al. 2022; Gustafsson et al. 2023) and improvements in sequencing performance (McCombie et al. 2019). It is desirable to build a simulation system for RNA-Seq because the characteristics of measurement errors differ between microarrays and RNA-Seq (Zhang et al. 2020). Specifically, it will be possible to easily extend the simulation system in this study by using the precision weights of voom (Law et al. 2014; Iwayama et al. 2017; Kashima et al. 2021). Although some of the results obtained from the simulation system for microarrays were successfully verified using actual RNA-Seq data (Fig. 7c, d), it would be beneficial to conduct more detailed verification using large-scale RNA-Seq data and a simulation system for RNA-Seq in the future.

The simulation system developed in this study and the findings obtained through the analysis will be useful in various aspects of field transcriptome research. For example, it can be used to generate synthetic data for the development of new modeling methods and for performance comparisons. It will also be useful for developing efficient sampling strategies to build more accurate models with fewer samples. Current field transcriptome modeling requires a large amount of data. If the amount of data required can be reduced, it will be possible to model a wider variety of species and strains. It is expected that the modeling of field transcriptomes in more species will reveal differences and similarities between species. Comparing field transcriptome models will be an important approach for understanding how organisms adapt to complex environments.

## Acknowledgements

We thank Dynacom Co., Ltd. (Chiba, Japan) for technical assistance. This study was supported by JST CREST Grant Number JPMJCR15O2, JST FOREST Grant Number JPMJFR210B, JSPS (JP20H00423, JP23H00386 and JP23K18156) and MEXT (JP23H04967) awarded to AJN.

## Author contributions

DE, YH, TM and AJN analyzed the data. AJN and DE conceived the study. DE and AJN wrote the manuscript with input from all co-authors.

## Data availability

The scripts used in this study are available at https://github.com/naganolab/Sim.factors.transcriptome_model

---

**Supplementary Figure S1.**
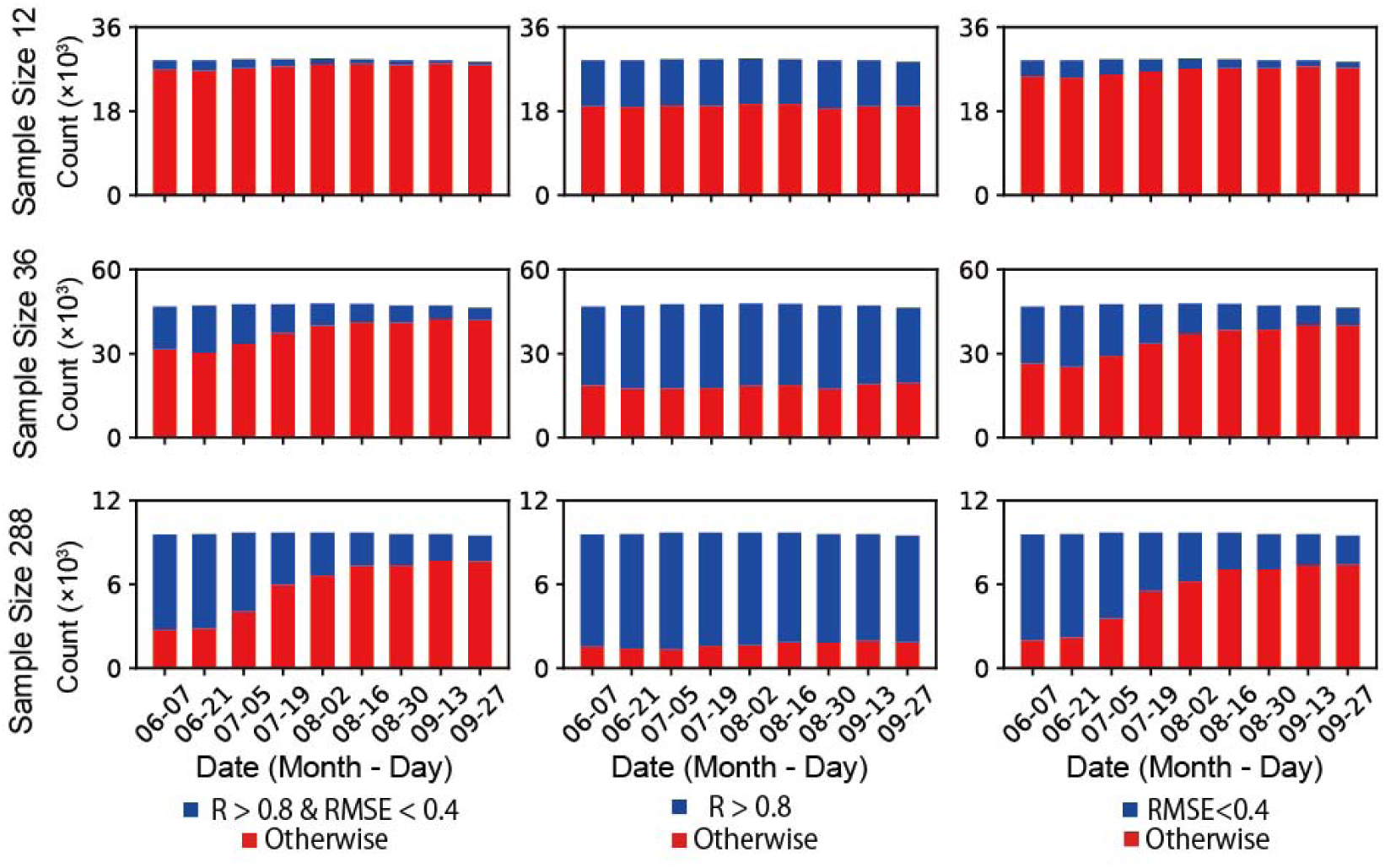
Counts of data exceeding the threshold for the correlation coefficient (R) and RMSE in sample size 12, 36 and 288. The correlation coefficient and RMSE were calculated for each gene, each simulation sampling condition, and each day of the evaluation data.

**Supplementary Figure S2.**
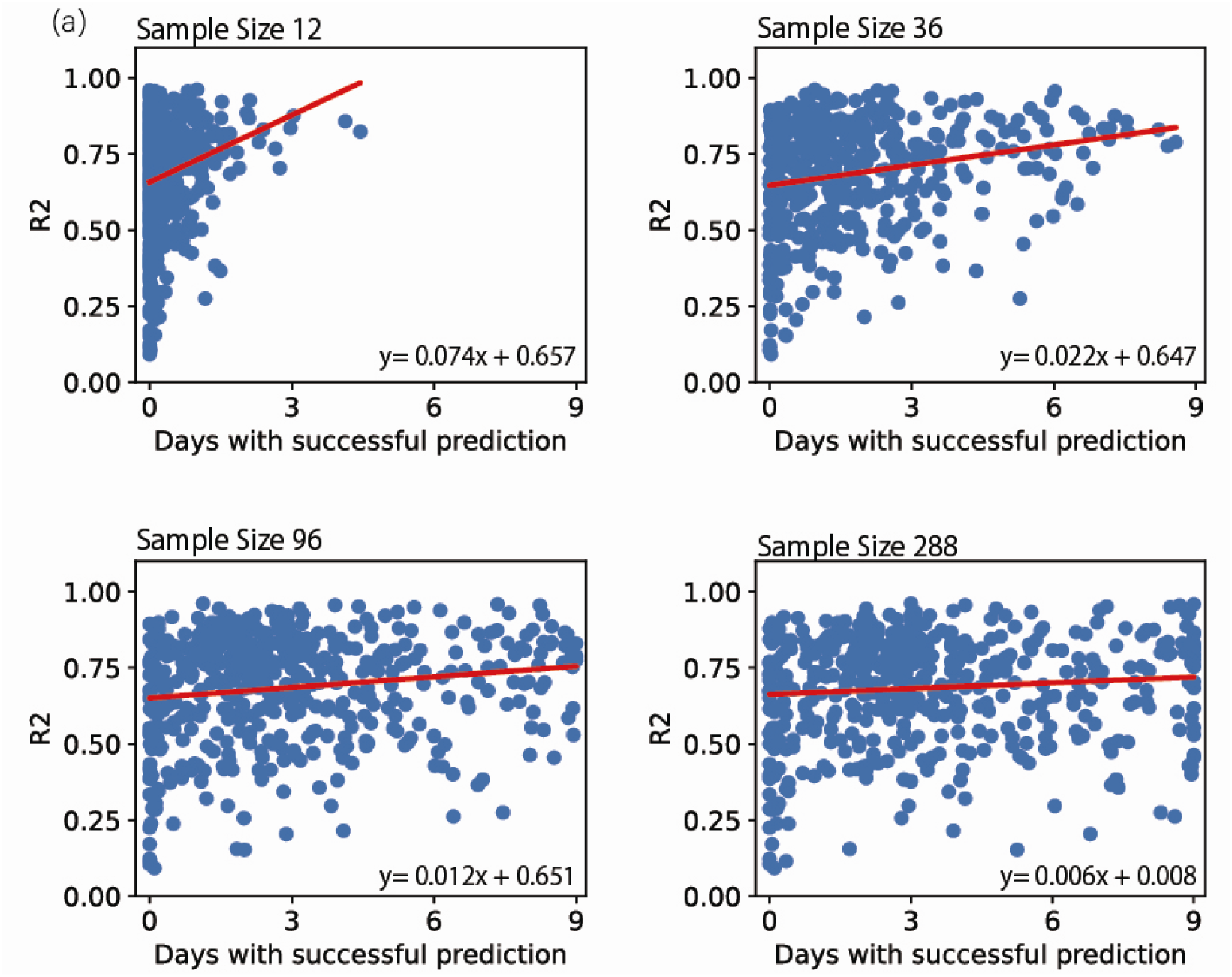
Effects of the relative amplitudes of gene expression noise to the deterministic expression dynamics on prediction accuracy. Vertical axis indicates the coefficients of determination of the reference FIT model for training data: 461 samples of microarray data. Horizontal axis indicates the average number of days of successful prediction of the test FIT models.

**Supplementary Figure S3.**
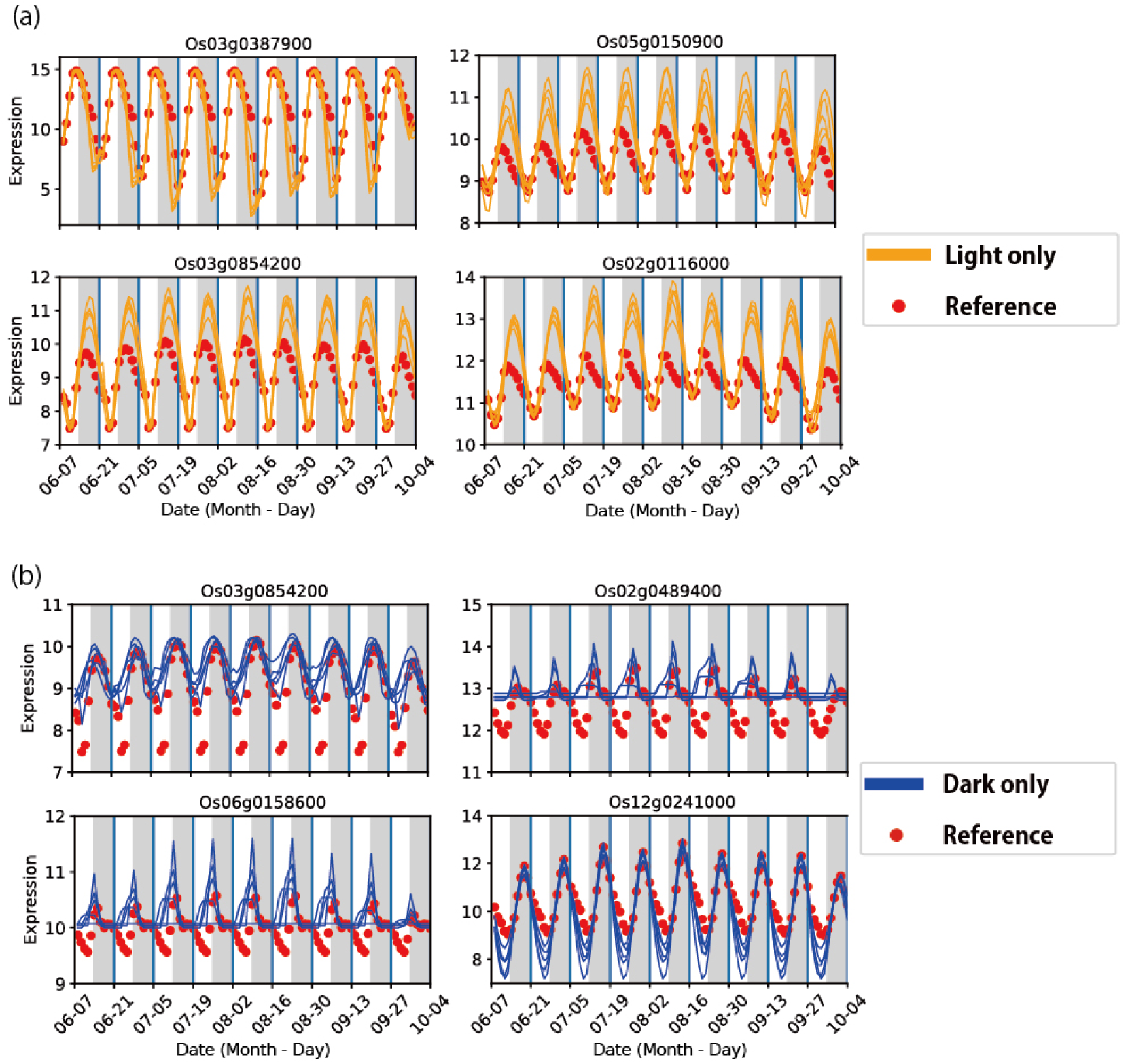
Temporal expression patterns of genes with low prediction accuracy in light-only/dark-only conditions. (a, b) Examples of genes with successful prediction in 9 days of evaluation data in the control condition and in no day in the light-only (a) or dark-only (b) condition. Orange and blue lines indicate predicted gene expressions of the test FIT models in 5 simulation sampling conditions. Red points indicate expected gene expressions of the reference FIT model.

**Supplementary Figure S4.**
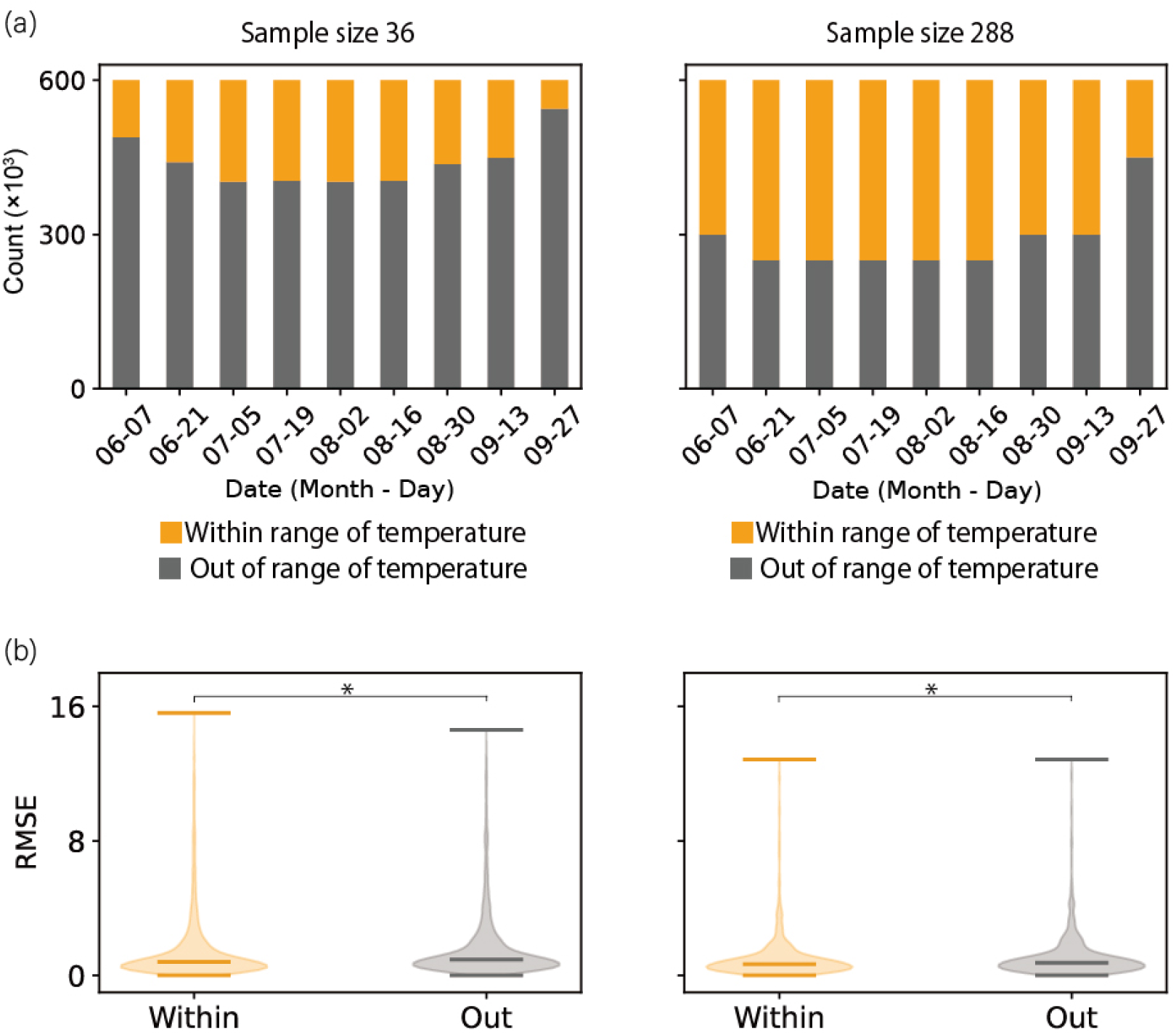
Prediction accuracy in the unsampled temperature range in simulation sampling conditions with 36 and 288 samples. (a) Data counts temperature of the evaluation data within (orange) and out of (gray) the range of training data. (b) Prediction error within and out of the temperature range of the training data, for simulation sampling conditions with 36 samples (left) and 288 samples (right). *: P < 0.001

**Supplementary Figure S5.**
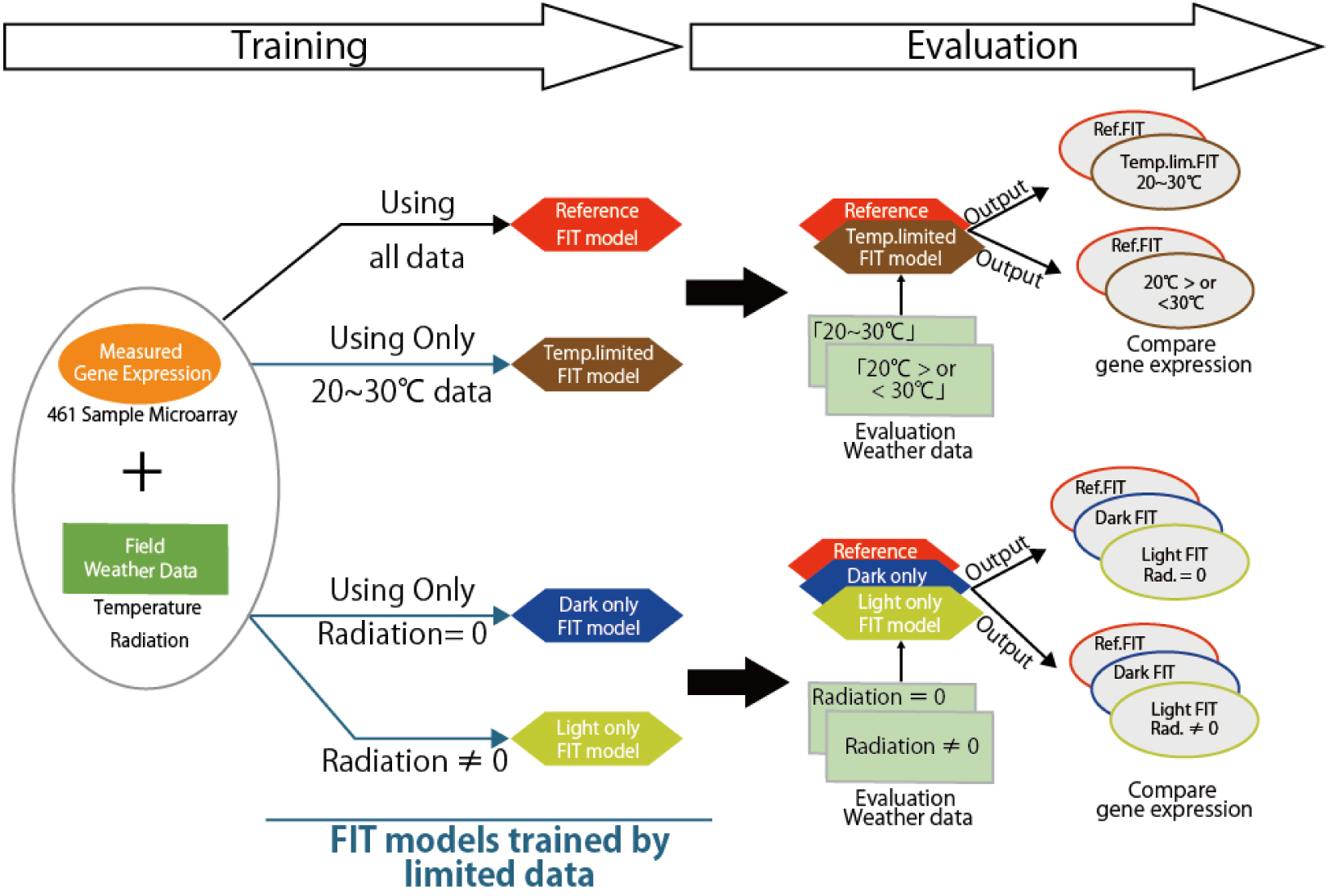
Training and evaluation of the FIT models trained by subsets of training data for the reference FIT model. The circles indicate gene expression data. The hexagons indicate FIT models. The squares indicate weather data.

